# Metabolomic signature of the maternal microbiota in the fetus

**DOI:** 10.1101/2020.09.28.317081

**Authors:** Tiina Pessa-Morikawa, Aleksi Husso, Olli Kärkkäinen, Ville Koistinen, Kati Hanhineva, Antti Iivanainen, Mikael Niku

## Abstract

The maternal microbiota affects the development of the offspring by microbial metabolites translocating to the fetus. We investigated samples of placenta, fetal intestine and brain from germ-free (GF) and specific pathogen free (SPF) mouse dams by non-targeted metabolic profiling. One hundred one annotated metabolites and altogether 3680 molecular features were present in significantly different amounts in the placenta and/or fetal organs of GF and SPF mice. The concentrations of more than half of the annotated and differentially expressed metabolites were lower in the GF organs, suggesting their microbial origin or a metabolic response of the host to the presence of gut microbiota. The clearest separation was observed in the placenta. Metabolites that were detected in lower amounts in the fetal organs in the GF mice included 5-aminovaleric acid betaine, trimethylamine N-oxide, catechol-O-sulphate, hippuric and pipecolic acid. Derivatives of the amino acid tryptophan, such as kynurenine, 3-indolepropionic acid and hydroxyindoleacetic acid, were also decreased in the absence of microbiota. Several metabolites had higher levels in the GF mice. These could be precursors of microbial metabolites or indicators of host metabolic response to the absence of gut microbiota. Ninety-nine molecular features were only detected in the SPF mice, suggesting the existence of yet unidentified microbially modified metabolites that potentially influence fetal development.

## 1 Introduction

The intestinal microbiota has a great impact on the life and wellbeing of the host. Microbes residing in the gut participate in digestion and metabolic modification of nutrients, producing substances that are absorbed by the host (Zhang and Davies, 2016). The cell numbers of gut microbiota are estimated to at least equal and its gene pool exceed that of its host (Sender et al., 2016; Lloyd-Price et al., 2017). The potential of the microbiota to influence host metabolism is illustrated by a comparison of the serum metabolome of conventionally colonized and germ-free (GF) mice: 3.5% of the > 4000 molecular features detected were unique for conventional mice and 10% of the shared molecular features had significantly different levels between the groups (Wikoff et al., 2009). All the organ systems of the host are affected to a varying degree (Quinn et al., 2020).

While a majority of the compounds originating from microbial metabolism detected in mammalian tissues still remain uncharacterized, some of these substances and their effects on the host are well documented. These include short chain fatty acids (SCFAs) which the host utilizes as an essential part of its metabolism (LeBlanc et al., 2017). The SCFAs produced from complex carbohydrates by microbes residing in the alimentary tract are an important source of energy for the host (Bergman, 1990). The SCFAs have also been shown to contribute to the maintenance of the gut epithelium and regulation of the immune responses by facilitating regulatory T cell generation in the colonic mucosa (Roediger and Moore, 1981; Inan et al., 2000; Arpaia et al., 2013; Furusawa et al., 2013; Smith et al., 2013). Gut-residing microbes are also known to modify endogenous primary bile acids creating molecular species such as deoxycholate that stimulates colonic enteroendocrine cells and thus affects the regulation of the intestinal function of the host (Yano et al., 2015). Other microbial metabolites or their host-produced derivatives, such as trimethylamine N-oxide (TMAO) and 5-aminovaleric acid betaine (5-AVAB), are known to modify specific host reactions of lipid metabolism (Wang et al., 2011; Koeth et al., 2013; Kärkkäinen et al., 2018). Microbiota also affects the levels of intestinal and absorbed nutrients, particularly amino acids (Wikoff et al., 2009; Mardinoglu et al., 2015; Yamamoto et al., 2018). Microbial metabolites of amino acids, such as the indole derivatives of tryptophan, are signaling molecules with both local and systemic effects (Agus et al., 2018).

The metabolic coexistence between the animal and the bacteria begins already before birth. While it is still unclear whether small numbers of live microbes exist in the healthy fetus, hundreds of microbial metabolites originating from the dam pass through the placenta (Gomez de Aguero et al., 2016; Walker et al., 2017). Very little is known of their properties and physiological effects across developing organs of the fetus (Ganal-Vonarburg et al., 2020). Microbe-derived aryl hydrocarbon receptor (AhR) ligands and microbially regulated retinoids are essential for fetal development of the immune system (van de Pavert et al., 2014; Gomez de Aguero et al., 2016; Grizotte-Lake et al., 2018). Maternal SCFAs are also readily transmitted to the fetus, programming the fetal metabolic and neural systems (Kimura et al., 2020). Other maternally derived microbial metabolites have been primarily studied in the context of toxicology (Ganal-Vonarburg et al., 2020). These observations suggest that whole bacteria are not necessarily required to inflict inflammatory immune responses by the host cells (Horn et al., 2000).

To examine the extent of the cross-placental transfer of microbial metabolites during pregnancy, we analyzed placenta, fetal intestine and brain samples from germ-free and specific pathogen free (SPF) murine dams using a broad non-targeted metabolomics approach (Fig. 1). Ultra-high performance liquid chromatography (UHPLC) coupled with quadrupole time-of-flight (QTOF) mass spectrometry allowed the detection of thousands of differentially abundant molecular features in the organ samples.

**Figure 1.**
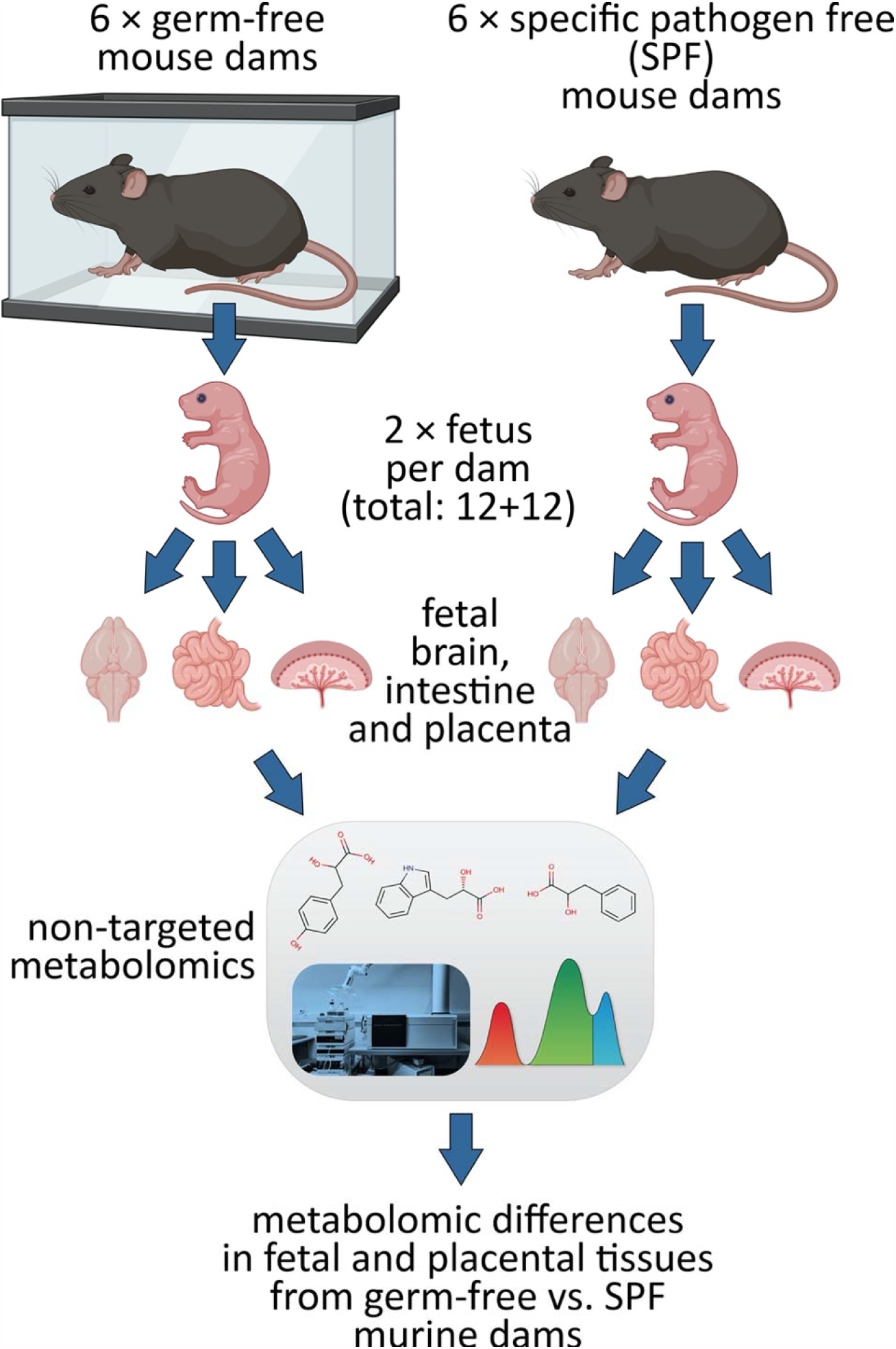
Experimental design. Fetal and placental tissues were collected from germ-free (GF) and specific pathogen free (SPF) mouse dams. The tissues were compared using non-targeted metabolomics, revealing metabolomic differences induced by maternal microbiological status in the fetuses. Figure created with Biorender.com.

## 2 Materials & Methods

### 2.1 Fetal and placental mouse organ samples

Fetal and placental mouse organ samples from pregnant germ-free (GF, n=6) and specific pathogen free (SPF, n=6) C57BL/6J dams were obtained from the EMMA Axenic Service at Instituto Gulbenkian de Ciência, Portugal. The dams were euthanized 18.5 days post coitum. Whole brain, intestine and placenta were collected from two fetuses per dam, to a total of 12 fetuses per experimental group. The fetal organ samples were immediately frozen in liquid nitrogen, stored at -80 °C and transferred on dry ice to the research laboratory. In both groups, there were 7 male fetuses and 5 female fetuses. The GF and SPF statuses were regularly monitored by culture and 16S qPCR. The GF dams were 3-4 months old and the SPF dams 4-5 months old. All dams were fed identical RM3-A-P breeding diets (SDS Special Diet Services, Essex, UK), autoclaved at 121 °C. The SPF feed was autoclaved for 20 minutes and the GF feed for 30 minutes due to logistical reasons.

### 2.2 Sample processing

Metabolomics analyses were performed at Afekta Technologies Ltd. Frozen organ samples were thawed at 8 °C for two hours and then weighed (approx. 100 mg) in homogenizer tubes. For the metabolite extraction, cold methanol (80 % v/v) was added in a ratio of 500 µl per 100 mg of sample. The samples were homogenized (TissueLyser II bead mill, Qiagen, Hilden, Germany) using metal beads at 6 m/s for 30 seconds. The samples were then shaken for 5 minutes in room temperature and centrifuged at 14,000 rpm at 4 °C for 10 min. After the centrifugation, the samples were kept on ice for 5 to 10 min, after which the supernatant was filtered (Acrodisc 0.2 µm PTFE membrane, Pall, Port Washington, NY, USA) into HPLC vials for analysis. The pooled quality control (QC) sample was prepared by collecting 20 µl from each sample vial and combining the material to two vials.

### 2.3 LC–MS analysis

The samples were analyzed by liquid chromatography–mass spectrometry, consisting of a 1290 Infinity Binary UPLC coupled with a 6540 UHD Accurate-Mass Q-TOF (Agilent Technologies Inc., Santa Clara, CA, USA), as described previously (Klåvus et al., 2020). In brief, a Zorbax Eclipse XDB-C18 column (2.1 × 100 mm, 1.8 µm; Agilent Technologies) was used for the reversed-phase (RP) separation and an Acquity UPLC BEH amide column (Waters Corporation, Milford, MA, USA) for the HILIC separation. After each chromatographic run, the ionization was carried out using jet stream electrospray ionization (ESI) in the positive and negative mode, yielding four data files per sample. The collision energies for the MS/MS analysis were selected as 10, 20 and 40 V, for compatibility with spectral databases.

### 2.4 Data analysis

Peak detection and alignment were performed in MS-DIAL ver. 4.00 (Tsugawa et al., 2015). For the peak collection, *m/z* values between 50 and 1500 and all retention times were considered. The amplitude of minimum peak height was set at 2000. The peaks were detected using the linear weighted moving average algorithm. For the alignment of the peaks across samples, the retention time tolerance was 0.05 min and the *m/z* tolerance was 0.015 Da. The heatmaps were produced with Multiple Experiment Viewer (MeV) version 4.9.0. MetaboAnalyst 4.0 was used for the pathway analysis of the annotated metabolites (Chong et al., 2018). Data clean-up (for each mode separately) and statistics (for all signals remaining after clean-up) were performed in R version 3.5.1. Molecular features were only kept if they met all the following quality metrics criteria: low number of missing values, present in more than 70% of the QC samples, present in at least 60% of samples in at least one study group, RSD* (the non-parametric version of relative standard deviation) below 20%, D-ratio* (non-parametric measure of the spread of the QC samples compared to the biological samples) below 10%. In addition, if either RSD* or D-ratio* was above the threshold, the features were still kept if their classic RSD, RSD* and basic D-ratio were all below 10%. Low-quality features were flagged and discarded from statistical analyses. Drift correction was applied to the data.

The cleaned data matrices of the four modes were combined before imputation. Features were then imputed using random forest imputation with an OOB error of 0.009. QC samples were removed prior to imputation to prevent them from biasing the procedure. Differential features between the treatment (GF) and control (SPF) were determined using a simple linear model (Student’s *t*-test) fit separately for each feature. The results were adjusted for multiple comparisons using Benjamini–Hochberg false discovery rate (FDR). FDR-adjusted *p* values (*q* values) below 0.05 were considered significant.

For the MS Peaks to Pathways analysis in MetaboAnalyst 4.0 (Chong et al., 2018), the data was first normalized by medians, cube root transformed, automatically scaled, and parametric statistical significances calculated with equal variances and adjusted *p* value (FDR) cutoff 0.05. In Peaks to Pathways, the molecular weight tolerance was set to 10 ppm, primary ions enforced, and adducts set based on the experimental data. Gene Set Enrichment analysis (Subramanian et al., 2005) and Mummichog version 1.0.10 (Li et al., 2013) were used.

### 2.5 Compound identification

The chromatographic and mass spectrometric characteristics (retention time, exact mass, and MS/MS spectra) of the significantly differential molecular features were compared with entries in an in-house standard library and publicly available databases, such as METLIN and HMDB, as well as with published literature. The annotation of each metabolite and the level of identification was given based on the recommendations published by the Chemical Analysis Working Group (CAWG) Metabolomics Standards Initiative (MSI): level 1 refers to confirmed identifications based on reference standards analyzed with the same instrument with identical conditions; level 2 means putative annotations with matching *m/z* and MS/MS fragmentation spectra with publicly available databases; level 3 signifies a putative characterization of compound class based on the observed physicochemical characteristics of the molecular feature; and level 4 covers all the remaining (unknown) signals (Sumner et al., 2007). Level 2 annotation was also given if MS/MS data was not available but the retention time and calculated molecular formula matched with that of a reference standard.

## 3 Results

### 3.1 Differences in all observed molecular features

The non-targeted metabolomics data consisted of a total of 12166 molecular features from four analytical modes after data cleanup. The metabolic profiles were clearly different in all studied organs (Fig. 2 and Table S1). The GF and SPF mice clustered separately in t-distributed stochastic neighbor embedding (TSNE) analysis, especially when each organ was analyzed individually (Fig. 2). The clearest separation between GF and SPF samples was observed in the placenta. The gender of the fetus did not influence the separation (not shown).

**Figure 2.**
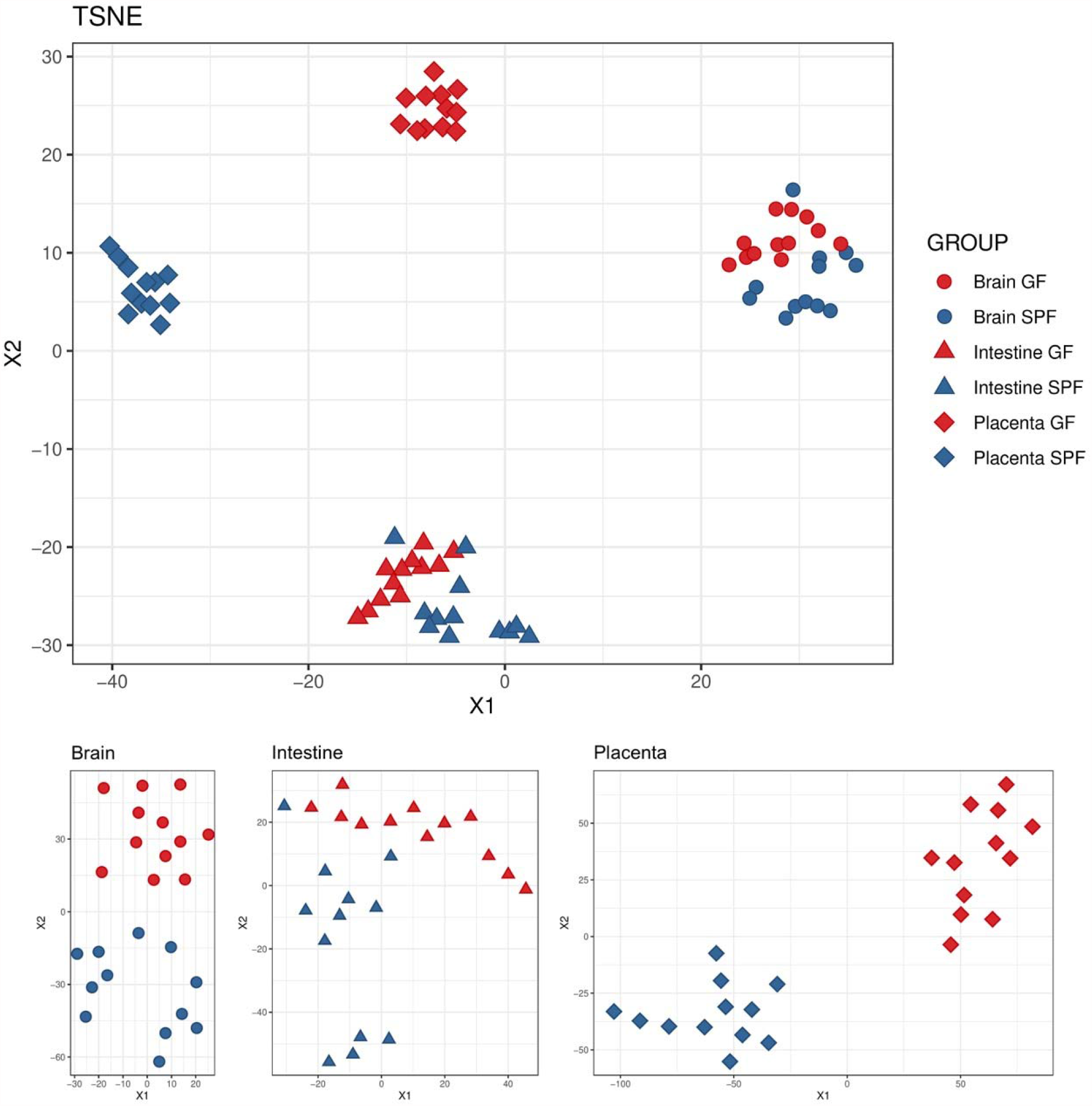
Metabolic profiles of germ-free (GF, red) and specific pathogen free (SPF, blue) placentae and fetal organs, analyzed by t-Distributed stochastic neighbor embedding (TSNE). The results are shown for the whole dataset including all the signals after data cleanup (n = 12 166) and separately for each organ.

The clustering by organ is also evident in the heatmap of all observed molecular features (Fig. 3). At this level, the difference in signal abundance related to germ-free status can be observed from a few relatively small clusters of molecular features in each analyzed organ.

**Figure 3.**
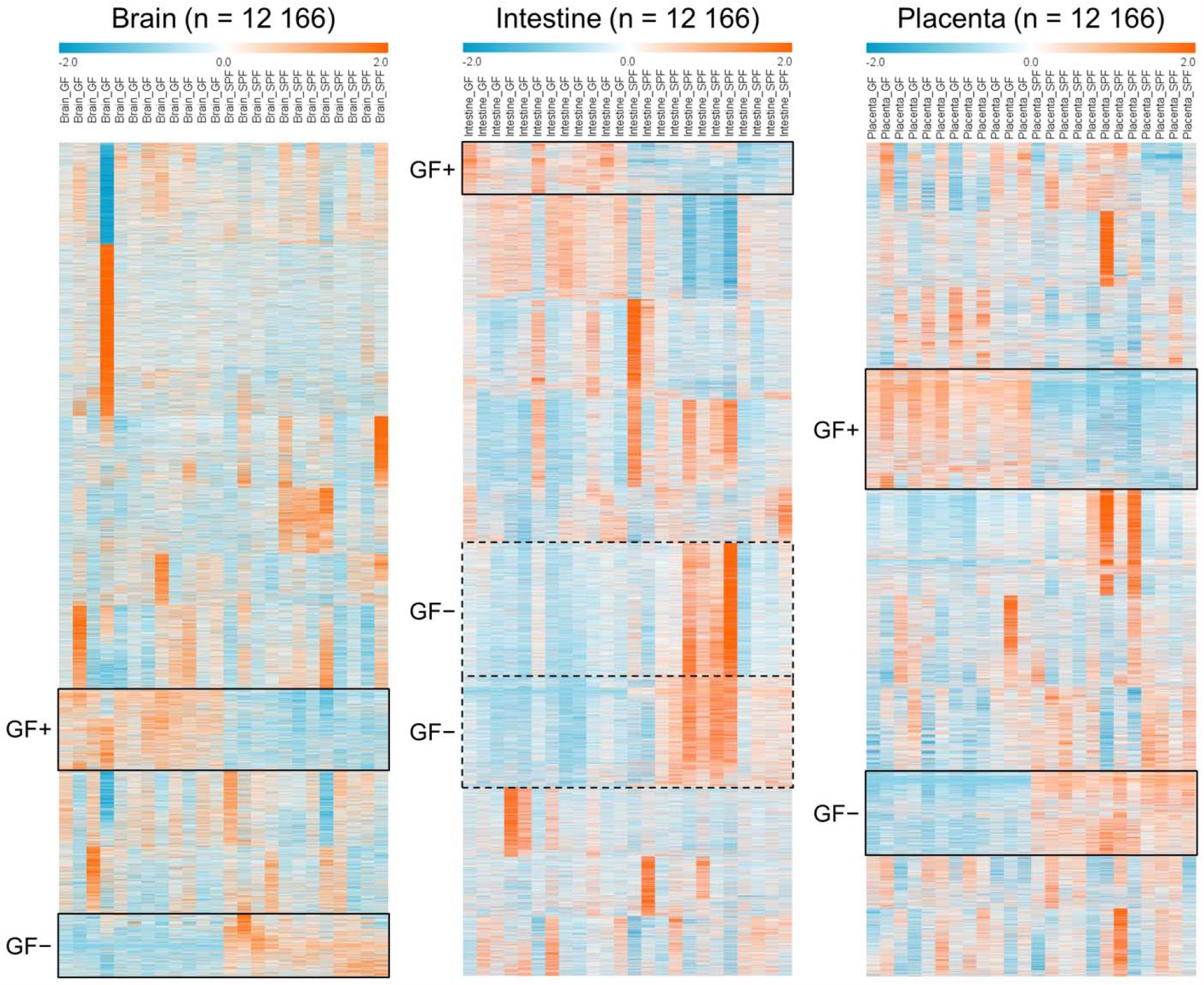
Heat map of the normalized abundances of all the signals from the four studied modes (RP+, RP−, HILIC+, HILIC−) after data cleanup (n = 12 166) in the brain, intestinal and placental samples. k-Means clustering (k = 10) was applied to each sample type separately to arrange the metabolites based on their similarity of the abundance between the samples. Clusters with tendency for increased or decreased abundance in the GF (germ-free) mice compared to SPF (specific pathogen free) mice are highlighted (continuous line: consistent trend across all samples, dashed line: trend with high interindividual variability). Individual signals are not aligned throughout all organ samples due to separately performed clustering.

The concentrations of 3680 molecular features differed between germ-free (GF) and specific pathogen free (SPF) mice in at least one of the organs investigated (FDR-adjusted *p* < 0.05 and Cohen’s *d* > 0.8). There were 2200 features which were more abundant in SPF mice in at least one organ (Fig. 4a). These were most numerous in the fetal intestine. One hundred sixty-eight features were more abundant in SPF mice in all three organs investigated. Similarly, 1533 features were more abundant in GF mice in at least one organ, most commonly in placenta (Fig. 4b). Eighty-eight features were more abundant in GF mice in all organs.

**Figure 4.**
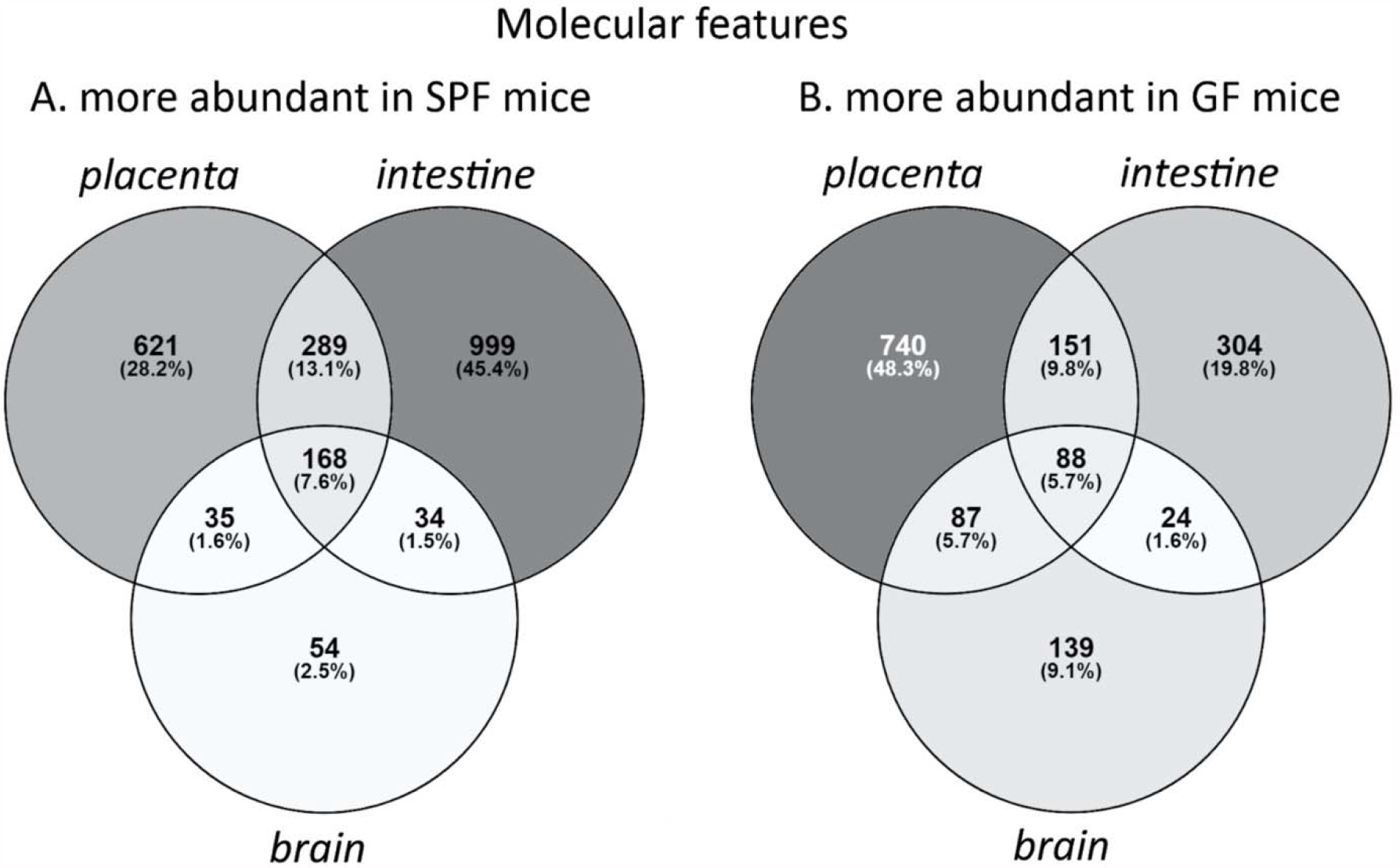
Venn diagrams showing the distribution of molecular features (n = 3680) which were more abundant (FDR-adjusted p < 0.05; Cohen’s d < -0.8 or > 0.8) A) in specific pathogen free mice in at least one organ, B) in germ-free mice in at least one organ. SPF = specific pathogen free, GF = germ-free.

A total of 99 features were only observed in SPF mice (Fig. 5). These were most commonly detected in all three SPF organs (n = 37), or in both placenta and fetal intestine (n = 36). None were detected only in both fetal organs, or only in the fetal brain. In contrast, only 6 features were observed exclusively in GF mice (not shown).

**Figure 5.**
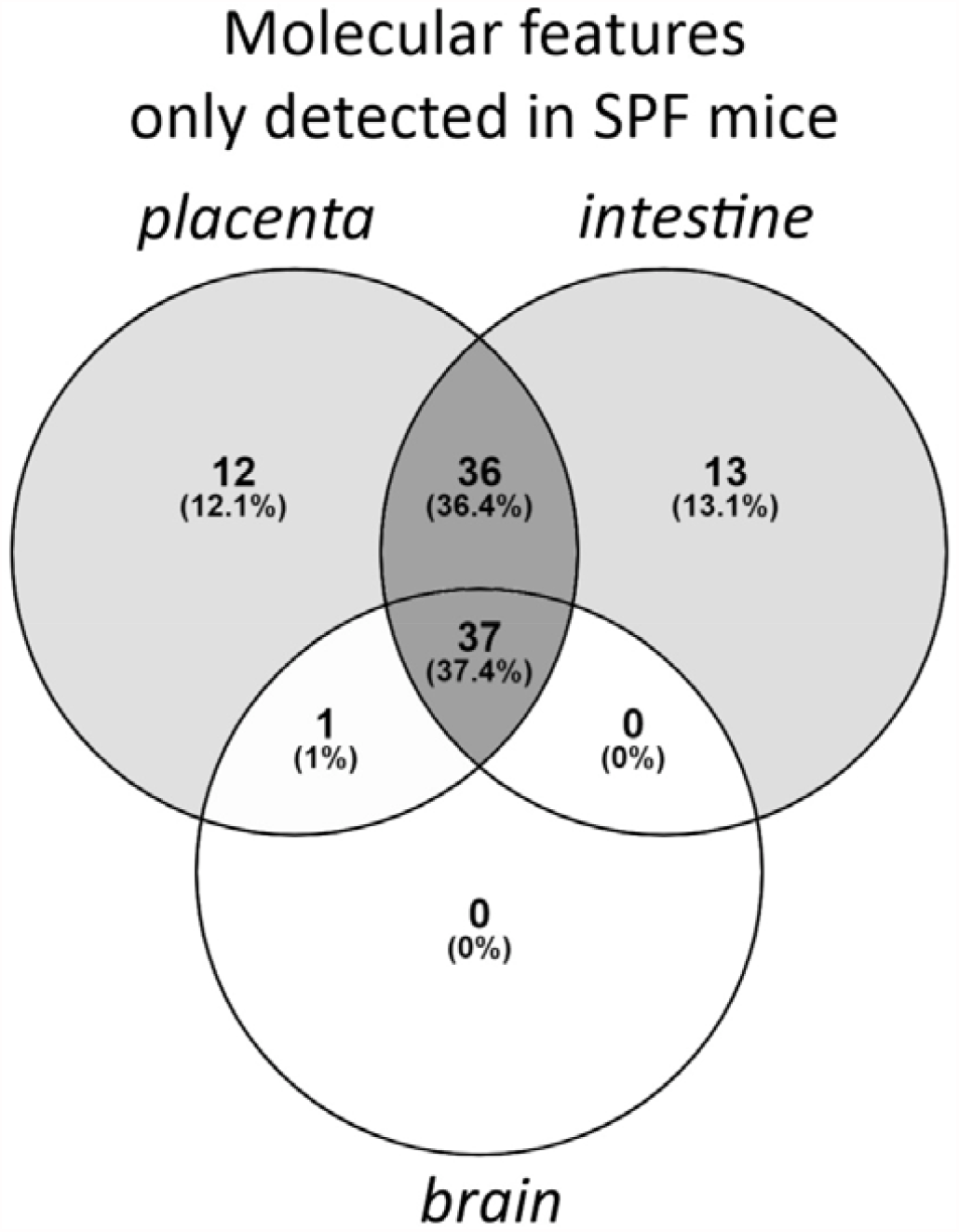
Venn diagram showing the distribution of molecular features (n = 99) which were observed only in specific pathogen free mice (signal-noise ratio > 5 in SPF mice and < 5 in GF mice; FDR-adjusted p < 0.05, Cohen’s d < -0.8 for GF vs. SPF). SPF = specific pathogen free, GF = germ-free.

### 3.2 Annotated metabolites

Among the differentially abundant molecular features, 101 metabolites were annotated. Out of these, 54 were identified (MSI level 1), 35 putatively annotated (level 2), 10 putatively characterized for compound class (level 3), and 2 unknowns (level 4) given a molecular formula (Table S1). A heatmap of significantly differential annotated metabolites is shown in Figure 6. Sixty-one of these metabolites were more abundant in SPF mice in at least one organ, most of these in all three organs or in intestine and/or placenta (FDR-adjusted *p* < 0.05, Cohen’s *d* > 0.8; Fig. 6 & 8, Table S1).

**Figure 6.**
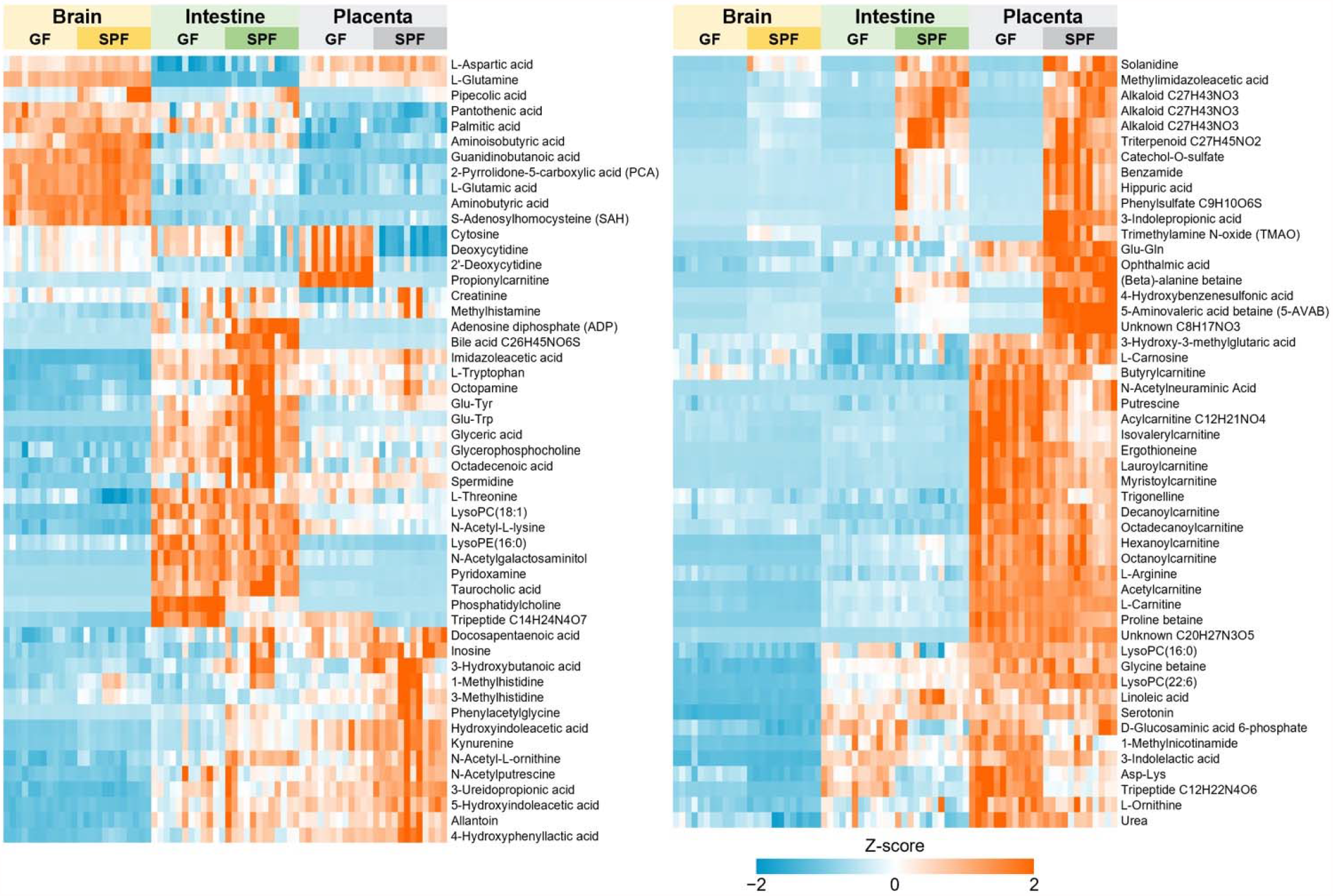
Heat map of the normalized signal abundances of the significantly differential annotated metabolites (n = 101) in each studied sample. A hierarchical clustering was applied to arrange the metabolites based on their similarity of the abundance between the samples.

Volcano plots of FDR-adjusted *p* values and Cohen’s *d* values for each organ are shown in Figure 7. Twenty-five metabolites were more abundant in all three SPF organs. Of these, 5-AVAB [also known as δ-valerobetaine (δVB), N,N,N-trimethyl-5-aminovalerate (TMAV) and N,N,N-trimethyl-5-aminovaleric acid (TMAVA)] was the most affected metabolite in all three organs. Other metabolites significantly affected included TMAO, alanine / β-alanine betaine, solanidine, catechol-O-sulphate, hippuric and pipecolic acid, amino acids and their derivatives (such as kynurenine, 3-indolepropionic acid and aminoisobutyric acid) and some small peptides. Five of the annotated compounds were observed exclusively in SPF mice: benzamide, 4-hydroxybenzenesulfonic acid, two unidentified alkaloids and a triterpenoid. Forty annotated metabolites were more abundant in GF mice in at least one organ, primarily in placenta and/or brain (Fig 8). These included several acylcarnitines, phosphatidylcholine, amino acids (such as L-threonine, L-arginine, ergothioneine and L-ornithine) and several small peptides.

**Figure 7.**
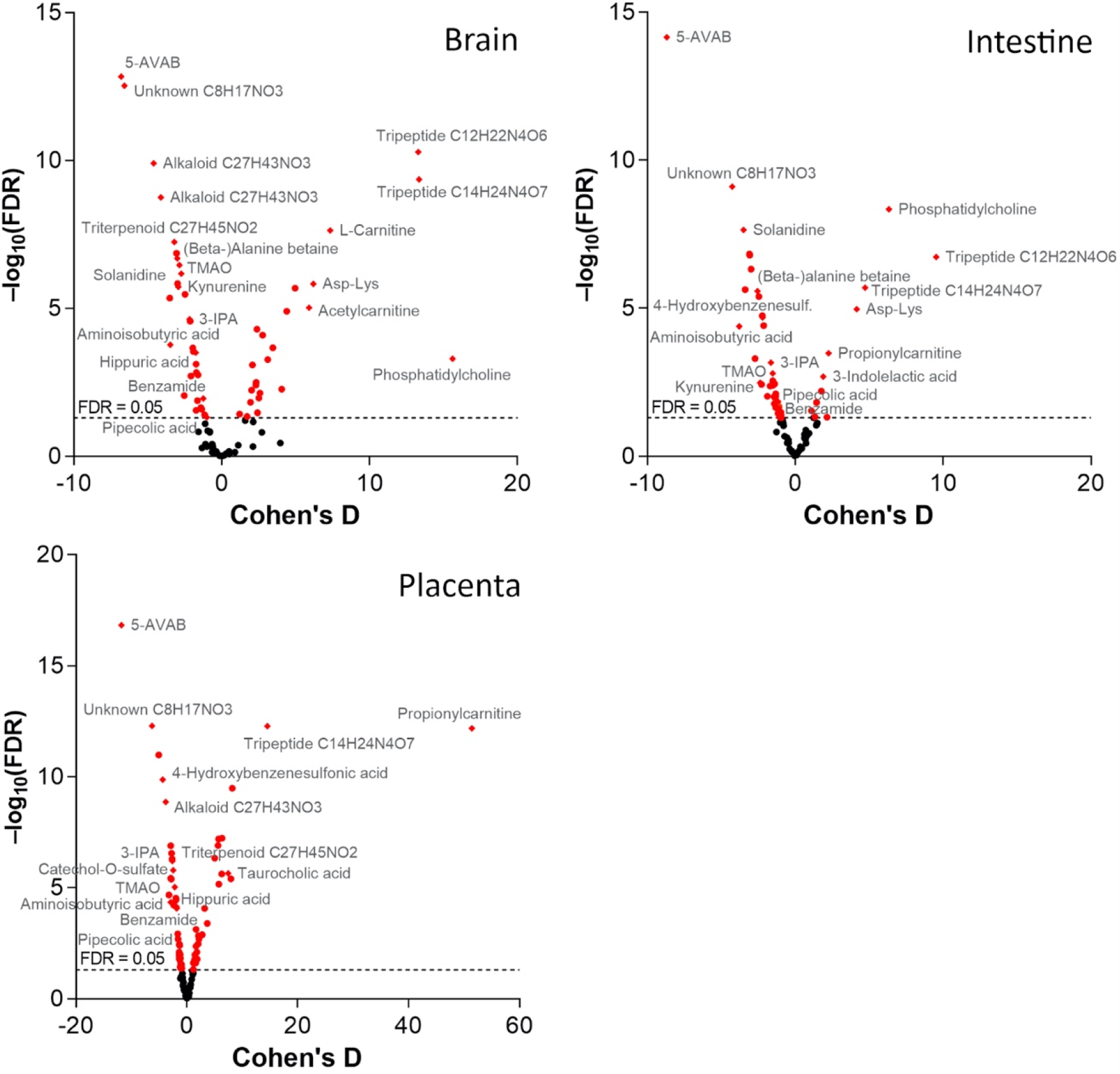
Volcano plots of the FDR-adjusted p values and Cohen’s d values for the significantly differential annotated metabolites (n = 101) in germ-free vs. specific pathogen free mice. Named compounds are represented as diamonds.

**Figure 8.**
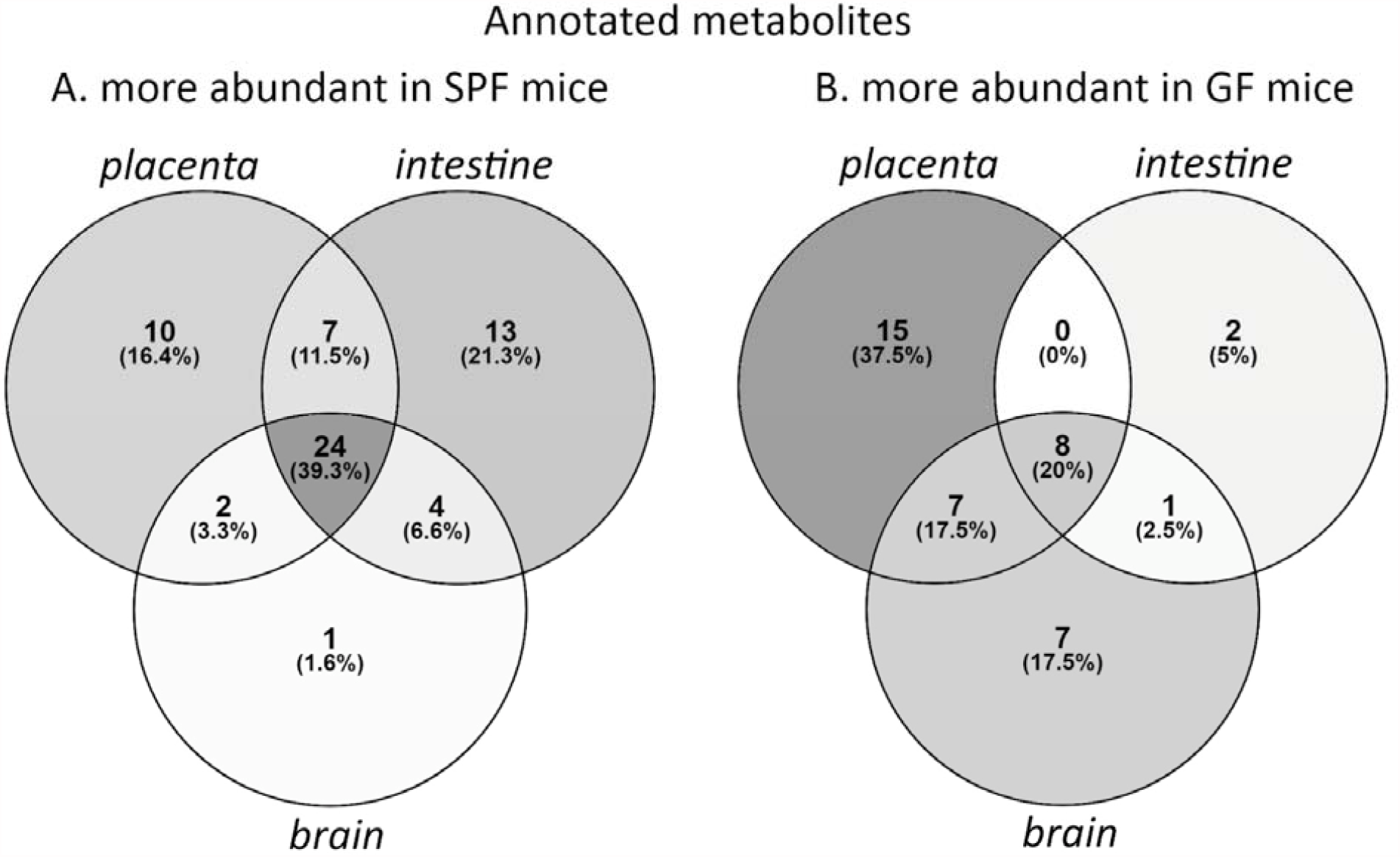
Venn diagrams showing the distribution of annotated metabolites which were more abundant (FDR-adjusted p < 0.05; Cohen’s d > 0.8) A) in specific pathogen free mice in at least one organ (n = 61), B) in germ-free mice in at least one organ (n = 40). SPF = specific pathogen free, GF = germ-free.

### 3.3 Pathway analysis

In order to predict metabolic pathways affected by the lack of microbiota in germ-free mice, we performed the MS Peaks to Pathways analysis in the MetaboAnalyst pipeline (Chong et al., 2018). This module utilizes the mummichog algorithm and gene set enrichment analysis (GSEA), which fit the mass spectrometry peak data into known metabolic pathways without pre-existing compound annotations (Chong et al., 2018).

In terms of murine KEGG and BIOCYC pathways, the metabolism of several essential amino acids, as well as tRNA charging / aminoacyl-tRNA biosynthesis were significantly different in both fetal organs (Supplementary Table 2) (Kanehisa et al., 2011; Karp et al., 2019). In the fetal intestine, phosphonate & phosphinate metabolism, glycolysis, caffeine metabolism, degradation of putrescine and nicotine, methionine salvage, and lipoate biosynthesis and incorporation were affected. In placenta, the affected pathways included lactose degradation, methionine salvage, nicotine degradation and folate biosynthesis.

## 4 Discussion

This is the first study probing the effects of a complete maternal microbiota on the metabolite composition in mammalian placental and fetal tissues. We compared fetuses of germ-free (GF) and specific pathogen free (SPF) murine dams utilizing non-targeted metabolomics. The lack of maternal microbiota affected the metabolite profile of the placenta, fetal intestine and brain. A total of 2200 detected molecular features were more abundant in SPF mice in at least one organ, while more than 1500 showed higher levels in the GF mice. Approximately one hundred molecular features could be detected only in the SPF organ samples. The numbers of compounds depleted in GF mice were largest in the fetal intestine and/or placenta. These observations indicate that the maternal microbiota strongly affects the host metabolism in placenta and in the fetus not only by directly producing metabolites but also pervasively impacting host physiology. One hundred one of the differentially abundant metabolites could be annotated based on current databases. Several betaines, amino acids and their derivatives, such as pipecolic acid and small peptides, certain alkaloids, catechol-O-sulphate and hippuric acid were more abundant in SPF mice. Several acylcarnitines, some amino acids, several small peptides and phosphatidylcholine were more abundant in GF mice.

### 4.1 Trimethylated compounds 5-AVAB, (β-)alanine betaine and TMAO

Considerably lower levels of the trimethylated componds, 5-AVAB, (β-)alanine betaine and TMAO, were detected in all the studied organs of the GF mice. These compounds are zwitterions containing a positively charged trimethylammonium group and a negatively charged oxygen. They can act as osmoprotective substances and methyl donors in various cellular metabolic processes (Raman and Rathinasabapathi, 2003, Velasquez et al., 2016; Koistinen et al., 2019). 5-AVAB is a bacterial metabolite (Servillo et al., 2018; Koistinen et al., 2019) with reported effects on fatty acid metabolism. It inhibits oxygen consumption due to β-oxidation of fatty acids in cultured mouse cardiomyocytes (Kärkkäinen et al., 2018). Thus, it may protect the heart tissue in ischemic conditions. The cord plasma of pre-eclamptic infants has been shown to contain increased levels of 5-AVAB (Jääskeläinen et al., 2018; Jääskeläinen et al., 2020). 5-AVAB has also been detected in the fetal mouse brain where its quantity has been shown to be strongly dependent on the presence of maternal gut microbiota (Vuong et al., 2020). Lower levels of (β-) alanine betaine have been detected in the intestine of GF mice than in conventional mice suggesting that it can also be produced by gut microbiota (Koistinen et al., 2019). Its role in mammalian physiology is poorly known.

TMAO is the mammalian end product of dietary phosphatidylcholine, choline, and carnitine, which are first metabolized by gut microbes into trimethylamine (TMA) and further into its *N*-oxide form in the liver (Al-Waiz et al., 1992; Lang et al., 1998). Whilst red meat is the main dietary source for the TMAO precursors choline and carnitine, TMAO as such is abundant in seafood (Landfald et al., 2017). TMAO inhibits reverse cholesterol transport by affecting bile acid synthesis on multiple levels and increases deposition of cholesterol to arterial walls (Koeth et al., 2013). Elevated levels are associated with increased risk of cardiovascular diseases, such as atherosclerosis and thrombosis (Tang et al., 2013; Zhu et al., 2016), but a causative role of TMAO has not been definitely established (Velasquez et al., 2016). A recent study by Vuong et al. (2020) indicates that TMAO and possibly also 5-AVAB have a role in development of the fetal brain. More than twofold reductions of TMAO and 5-AVAB were detected in the maternal blood and fetal brain of both GF and antibiotic-treated pregnant mice relative to SPF controls. Colonization of GF dams with ingenious spore-forming bacteria increased the levels of these compounds in the fetal brain. The maternal microbiota was shown to promote fetal thalamocortical axonogenesis and the effect could be reproduced by supplementing microbiota-depleted dams with TMAO (Vuong et al., 2020).

### 4.2 Amino acids and related metabolites

Concentrations of several amino acids and their derivatives were significantly different in the organs of the test groups. The pathway analysis based on unannotated MS peak data further substantiated the broad impacts of maternal microbiota on amino acid metabolism and tRNA charging especially regarding essential amino acids. Earlier studies have shown that the GF status is associated with differences in the levels of several amino acids in plasma and intestine (Wikoff et al., 2009; Mardinoglu et al., 2015; Yamamoto et al., 2018). Notably, microbial metabolism can diversify the fates of the amino acid tryptophan. On the one hand, microbes can produce an array of microbial indole metabolites, such as indole-3-propionic acid (IPA), in the gut lumen. On the other, they can also act indirectly by modifying the host metabolism of tryptophan to its neuromodulatory metabolite serotonin by enterochromaffin cells and to kynurenine by immune and epithelial cells (Clarke et al., 2012; Zelante et al., 2013; Reigstad et al., 2015; Yano et al., 2015). Adult GF mice have been reported to exhibit lower levels of serotonin in plasma and brain but higher levels of tryptophan in plasma than in conventional mice (Wikoff et al., 2009; Clarke et al., 2012).

We observed lower levels of tryptophan and its metabolites serotonin, hydroxyindoleacetic acid (HIAA), kynurenine and 3-indolepropionic acid (3-IPP) in intestine or brain or both of the organs of the GF fetuses. The levels of serotonin and 3-IPP were also reduced in placenta. Kynurenine and HIAA can act as ligands for AhR receptors expressed on gut epithelia and many types of immune cells and may have a significant role in modifying the host mucosal immune system to promote the survival of commensal microbiota and provide protection against pathogens (Zelante et al., 2013). Modulation of the levels of tryptophan, serotonin and kynurenine by microbiota may have effects on both the enteral and central nervous systems and the immune system. Disturbances in the intestinal microbiota can affect the levels of these metabolites, which may contribute to the pathogenesis of a multitude of disorders, such as inflammatory bowel diseases, metabolic syndrome and obesity, and various neuropsychiatric disorders (Agus et al., 2018). 3-indolepropionic acid is a strong antioxidant (Chyan et al., 1999; Hardeland et al., 1999). It has been shown to regulate the intestinal barrier function by acting as a ligand to the pregnane X receptor (PXR) (Venkatesh et al., 2014). It also has neuroprotective functions (Chyan et al., 1999) and was recently shown to promote thalamocortical axonogenesis in fetal mouse brain (Vuong et al., 2020). The changes in the levels of tryptophan metabolites in fetal intestine and brain reported here suggest that the microbial influences may begin already during fetal development.

Methylimidazoleacetic acid is the main metabolite of histamine. In this study, it was not detected in brain of the GF fetuses and its levels were also considerably lower in the other organs compared to SPF mice. Methylimidazoleacetic acid may be associated with miscarriage, potentially by the dysregulation of cytokine networks caused by imbalance in gut bacteria (Liu et al., 2020). Another main metabolic pathway of histamine leads to the formation of imidazoleacetic acid. Its levels were decreased in the placenta and fetal intestine of the GF mice. The 4 to 6-fold lower levels in GF mice fetal organs of pipecolic acid, a degradation product of L-lysine produced by intestinal bacteria, may reflect lack of the contribution of gut microbiota to the pipecolic acid pathway of lysine catabolism in GF mice (He, 2006). Other amino acid-related metabolites, including phenylacetylglycine, 1-methylhistidine, 3-methylhistidine, 3-hydroxybutanoic acid, Glu-Gln and Glu-Tyr, also had lower levels in the GF organs, indicating that the presence of microbiota may increase their production. Phenylacetylglycine has been identified as a gut microbial metabolite of phenylalanine with associations to health and disease in humans (Nemet et al., 2020). It has also been detected in forebrain of mice from neonatal period into adulthood (Swann et al., 2020). Certain amino acids and small peptides, such as L-threonine and two tripeptides with unknown structure, had higher levels in the GF organs compared to SPF, suggesting that they were accumulated in the organs due to lack of microbial metabolism.

### 4.3 Plant-derived metabolites catechol-O-sulfate, hippuric acid and solanidine

Catechol-*O*-sulfate and hippuric acid are metabolites of dietary polyphenols, such as flavonoids and phenolic acids, via degradation by gut microbiota and subsequent sulfation or glycine conjugation in the liver (Feliciano et al., 2016; de Mello et al., 2017). Hippuric acid is also produced from breakdown products of aromatic amino acids in liver, intestine and kidney for excretion into urine (Wikoff et al., 2009); gut microbiota is involved in this process (Lee et al., 2012). Levels of hippuric acid were significantly reduced in the placenta, fetal intestine and brain. Decreased levels of hippuric acid have been reported in serum of GF mice and in urine of antibiotic-treated rats (Wikoff et al., 2009; Lee et al., 2012) and recently also in the brain of fetuses of GF or antibiotic-treated mice (Vuong et al., 2020). Hippuric acid is one of the metabolites reported to be present in neonatal mouse brain with amounts decreasing by age (Swann et al., 2020). Apart from participating in removal of benzoate, a metabolite potentially toxic to mitochondria (Badenhorst et al., 2014), hippuric acid has been suggested a role in regulation of blood glucose levels, insulin secretion by β-cells and glucose utilization in skeletal muscle (de Mello et al., 2017; Bitner et al., 2018). Solanidine is a steroidal glycoalkaloid, which is obtained via dehydroxylation from other glycoalkaloids, such as α-chaconine and α-solanine, which are abundant in potato, a component of the RM3-A-P breeding diet. Solanidine has been detected as the main metabolite in rats after oral ingestion of α-chaconine, and the current findings support the hypothesis that solanidine is a gut microbial metabolite of dietary glycoalkaloids in mice (Norred et al., 1976).

### 4.4 Energy metabolism

The GF status is known to affect the energy metabolism of mice (Yu et al., 2019). We found increased levels of carnitine and various acylcarnitines in placenta and fetal organs of GF mice. These are precursors of TMAO (Velasquez et al., 2016), which suggests that they were not metabolized in the gut of the dam due to the lack of gut microbiota. Increased levels of carnitine could also be due to decreased levels of 5-AVAB in the GF mice, since 5-AVAB inhibits the cell membrane carnitine transporter responsible for cellular intake of carnitine (Kärkkäinen et al. 2018). Carnitine and acyl carnitines are involved in beta oxidation of fatty acids. They also have a multifaceted role in the developing brain (Ferreira and McKenna, 2017).

### 4.5 Unannotated molecular features undetected in GF mice

In total 3680 molecular features were differentially expressed in SPF and GF organs and 99 were not detected in GF mice. Most of these features could not yet be annotated. Unidentified metabolites strongly affected by the presence of maternal microbiota may contain hitherto unknown molecular species with potential bioactivity in the mammalian host, as was shown recently by identification of novel amino acid conjugations of bile acids by extensive bioinformatic analysis of metabolomics data from GF and SPF mice (Quinn et al., 2020).

### 4.6 Limitations of the study

The LC-MS method used to detect the metabolites in this study does not allow the detection of short-chain fatty acids (formic to valeric acid), many volatile organic compounds (VOCs) and molecules above 1500 Da (including lipopolysaccharides, large peptides, proteins, and nucleic acids). The SCFAs have been extensively studied already previously (Roediger and Moore, 1981; Inan et al., 2000; Smith et al., 2013; Reigstad et al., 2015; LeBlanc et al., 2017). SCFAs are known to cross the placenta to fetal tissues and have an impact on the fetal development (Shekhawat et al., 2003; Priyadarshini et al., 2014).

The current databases available for metabolite identification contain the most well-known mammalian endogenous metabolites. Reference data on exogenous (e.g. plant-derived) and microbially produced metabolites is limited. Therefore, a significant proportion of molecular features in the dataset could not be annotated. A tentative characterization of the chemical class was acquired for 10 of the unknowns with the prediction of molecular formula and comparison to *in silico* generated MS/MS fragmentation patterns.

### 4.8 Conclusion

The germ-free status of the dam strongly affects the metabolic landscape of the placenta and the developing fetus. Several known metabolites originating in microbial metabolism or combined metabolism of microbiota and the host were among the differential compounds, further highlighting the impact of gut microbiota on the host metabolism. Intriguingly, the most significantly differing metabolites included 5-AVAB and TMAO with reported effects on neural development in mice. Among these were also the tryptophan metabolite serotonin, an important regulator of neural and enteroendocrine functions, and kynurenine and HIAA that can act as effectors of the immune system through ArH receptors. Additionally, hundreds of unannotated molecular features were significantly more abundant in SPF mice or missing from GF mice, indicating that a majority of microbially processed metabolites in the fetus and placenta are still unknown. These represent a pool of compounds with potentially important bioactivities in the host and also effects on the fetal development of the brain and other organ systems.

## 6 Author contributions

MN, AI and KH designed the study. VK performed the metabolomics data processing and metabolite identification, OK and KH supervised the metabolomics analysis. All authors contributed to data analysis and interpretation. TPM, MN, VK and AH wrote the manuscript. All authors read, commented, and approved the final version of the manuscript.

## 8 Data availability

The metabolomics dataset can be found at EUDAT upon publication of the article.

## 9 Acknowledgements

We thank Kirsi Lahti and Santeri Suokas for expert technical assistance and Anton Klåvus for running the statistical analyses.

## References

Agus, A., Planchais, J., and Sokol, H. (2018). Gut Microbiota Regulation of Tryptophan Metabolism in Health and Disease. Cell Host Microbe 23, 716–724. doi:10.1016/j.chom.2018.05.003.

Al-Waiz, M., Mikov, M., Mitchell, S. C., and Smith, R. L. (1992). The exogenous origin of trimethylamine in the mouse. Metabolism 41, 135–136. doi:10.1016/0026-0495(92)90140-6.

Arpaia, N., Campbell, C., Fan, X., Dikiy, S., van der Veeken, J., deRoos, P., et al. (2013). Metabolites produced by commensal bacteria promote peripheral regulatory T-cell generation. Nature 504, 451–455. doi:10.1038/nature12726.

Badenhorst, C. P. S., Erasmus, E., van der Sluis, R., Nortje, C., and van Dijk, A. A. (2014). A new perspective on the importance of glycine conjugation in the metabolism of aromatic acids. Drug Metab. Rev. 46, 343–361. doi:10.3109/03602532.2014.908903.

Bergman, E. N. (1990). Energy contributions of volatile fatty acids from the gastrointestinal tract in various species. Physiol. Rev. 70, 567–590.

Bitner, B. F., Ray, J. D., Kener, K. B., Herring, J. A., Tueller, J. A., Johnson, D. K., et al. (2018). Common gut microbial metabolites of dietary flavonoids exert potent protective activities in β-cells and skeletal muscle cells. J. Nutr. Biochem. 62, 95–107. doi:10.1016/j.jnutbio.2018.09.004.

Chong, J., Soufan, O., Li, C., Caraus, I., Li, S., Bourque, G., et al. (2018). MetaboAnalyst 4.0: towards more transparent and integrative metabolomics analysis. Nucleic Acids Res. 46, W486–W494. doi:10.1093/nar/gky310.

Chyan, Y.-J., Poeggeler, B., Omar, R. A., Chain, D. G., Frangione, B., Ghiso, J., et al. (1999). Potent Neuroprotective Properties against the Alzheimer β-Amyloid by an Endogenous Melatonin-related Indole Structure, Indole-3-propionic Acid. J. Biol. Chem. 274, 21937– 21942. doi:10.1074/jbc.274.31.21937.

Clarke, G., McKernan, D. P., Gaszner, G., Quigley, E. M., Cryan, J. F., and Dinan, T. G. (2012). A distinct profile of tryptophan metabolism along the kynurenine pathway downstream of toll-like receptor activation in irritable bowel syndrome. Front. Pharmacol. 3, 90.

de Mello, V. D., Lankinen, M. A., Lindström, J., Puupponen-Pimiä, R., Laaksonen, D. E., Pihlajamäki, J., et al. (2017). Fasting serum hippuric acid is elevated after bilberry (Vaccinium myrtillus) consumption and associates with improvement of fasting glucose levels and insulin secretion in persons at high risk of developing type 2 diabetes. Mol. Nutr. Food Res. 61, 1700019. doi:10.1002/mnfr.201700019.

Feliciano, R. P., Boeres, A., Massacessi, L., Istas, G., Ventura, M. R., Nunes dos Santos, C., et al. (2016). Identification and quantification of novel cranberry-derived plasma and urinary (poly)phenols. Arch. Biochem. Biophys. 599, 31–41. doi:10.1016/j.abb.2016.01.014.

Ferreira, G. C., and McKenna, M. C. (2017). L-Carnitine and acetyl-L-carnitine roles and neuroprotection in developing brain. Neurochem. Res. 42, 1661–1675.

Furusawa, Y., Obata, Y., Fukuda, S., Endo, T. A., Nakato, G., Takahashi, D., et al. (2013). Commensal microbe-derived butyrate induces the differentiation of colonic regulatory T cells. Nature 504, 446–450. doi:10.1038/nature12721.

Ganal-Vonarburg, S. C., Hornef, M. W., and Macpherson, A. J. (2020). Microbial-host molecular exchange and its functional consequences in early mammalian life. Science 368, 604–607. doi:10.1126/science.aba0478.

Gomez de Aguero, M., Ganal-Vonarburg, S. C., Fuhrer, T., Rupp, S., Uchimura, Y., Li, H., et al. (2016). The maternal microbiota drives early postnatal innate immune development. Science 351, 1296–302. doi:10.1126/science.aad2571.

Grizotte-Lake, M., Zhong, G., Duncan, K., Kirkwood, J., Iyer, N., Smolenski, I., et al. (2018). Commensals Suppress Intestinal Epithelial Cell Retinoic Acid Synthesis to Regulate Interleukin-22 Activity and Prevent Microbial Dysbiosis. Immunity 49, 1103-1115.e6. doi:10.1016/j.immuni.2018.11.018.

Hardeland, R., Zsizsik, B. K., Poeggeler, B., Fuhrberg, B., Holst, S., and Coto-Montes, A. (1999). “Indole-3-Pyruvic and -Propionic Acids, Kynurenic Acid, and Related Metabolites as Luminophores and Free-Radical Scavengers,” in Tryptophan, Serotonin, and Melatonin, eds. G. Huether, W. Kochen, T. J. Simat, and H. Steinhart (Boston, MA: Springer US), 389–395. doi:10.1007/978-1-4615-4709-9_49.

He, M. (2006). Pipecolic acid in microbes: biosynthetic routes and enzymes. J. Ind. Microbiol. Biotechnol. 33, 401–407. doi:10.1007/s10295-006-0078-3.

Horn, D. L., Morrison, D. C., Opal, S. M., Silverstein, R., Visvanathan, K., and Zabriskie, J. B. (2000). What are the microbial components implicated in the pathogenesis of sepsis? Report on a symposium. Clin. Infect. Dis. 31, 851–858.

Inan, M. S., Rasoulpour, R. J., Yin, L., Hubbard, A. K., Rosenberg, D. W., and Giardina, C. (2000). The luminal short-chain fatty acid butyrate modulates NF-κB activity in a human colonic epithelial cell line. Gastroenterology 118, 724–734.

Jääskeläinen, T., Kärkkäinen, O., Jokkala, J., Litonius, K., Heinonen, S., Auriola, S., et al. (2018). A Non-Targeted LC-MS Profiling Reveals Elevated Levels of Carnitine Precursors and Trimethylated Compounds in the Cord Plasma of Pre-Eclamptic Infants. Sci. Rep. 8, 14616. doi:10.1038/s41598-018-32804-5.

Jääskeläinen T, Kärkkäinen O, Jokkala J, Klåvus A, Heinonen S, Auriola S, Lehtonen M, FINNPEC, Hanhineva K, Laivuori H. (2020). A non-targeted LC-MS metabolic profiling of pregnancy: longitudinal evidence from healthy and pre-eclamptic pregnancies. Metabolomics. Accepted manuscript in press.

Kanehisa, M., Goto, S., Sato, Y., Furumichi, M., and Tanabe, M. (2011). KEGG for integration and interpretation of large-scale molecular data sets. Nucleic Acids Res. 40, D109–D114. doi:10.1093/nar/gkr988.

Kärkkäinen, O., Tuomainen, T., Koistinen, V., Tuomainen, M., Leppänen, J., Laitinen, T., et al. (2018). Whole grain intake associated molecule 5-aminovaleric acid betaine decreases β-oxidation of fatty acids in mouse cardiomyocytes. Sci. Rep. 8, 13036. doi:10.1038/s41598-018-31484-5.

Karp, P. D., Billington, R., Caspi, R., Fulcher, C. A., Latendresse, M., Kothari, A., et al. (2019). The BioCyc collection of microbial genomes and metabolic pathways. Brief. Bioinform. 20, 1085–1093.

Kimura, I., Miyamoto, J., Ohue-Kitano, R., Watanabe, K., Yamada, T., Onuki, M., et al. (2020). Maternal gut microbiota in pregnancy influences offspring metabolic phenotype in mice. Science 367. doi:10.1126/science.aaw8429.

Klåvus, A., Kokla, M., Noerman, S., Koistinen, V. M., Tuomainen, M., Zarei, I., et al. (2020). “Notame”: Workflow for Non-Targeted LC–MS Metabolic Profiling. Metabolites 10, 135. doi:10.3390/metabo10040135.

Koeth, R. A., Wang, Z., Levison, B. S., Buffa, J. A., Org, E., Sheehy, B. T., et al. (2013). Intestinal microbiota metabolism of l-carnitine, a nutrient in red meat, promotes atherosclerosis. Nat. Med. 19, 576–585. doi:10.1038/nm.3145.

Koistinen, V. M., Kärkkäinen, O., Borewicz, K., Zarei, I., Jokkala, J., Micard, V., et al. (2019). Contribution of gut microbiota to metabolism of dietary glycine betaine in mice and in vitro colonic fermentation. Microbiome 7, 103. doi:10.1186/s40168-019-0718-2.

Landfald, B., Valeur, J., Berstad, A., and Raa, J. (2017). Microbial trimethylamine-N -oxide as a disease marker: something fishy? Microb. Ecol. Health Dis. 28, 1327309. doi:10.1080/16512235.2017.1327309.

Lang, D., Yeung, C., Peter, R., Ibarra, C., Gasser, R., Itagaki, K., et al. (1998). Isoform specificity of trimethylamine N-oxygenation by human flavin-containing monooxygenase (FMO) and P450 enzymes. Biochem. Pharmacol. 56, 1005–1012. doi:10.1016/S0006-2952(98)00218-4.

LeBlanc, J. G., Chain, F., Martín, R., Bermúdez-Humarán, L. G., Courau, S., and Langella, P. (2017). Beneficial effects on host energy metabolism of short-chain fatty acids and vitamins produced by commensal and probiotic bacteria. Microb. Cell Factories 16, 1– 10.

Lee, S. H., An, J. H., Park, H.-M., and Jung, B. H. (2012). Investigation of endogenous metabolic changes in the urine of pseudo germ-free rats using a metabolomic approach. J. Chromatogr. B 887–888, 8–18. doi:10.1016/j.jchromb.2011.12.030.

Li, S., Park, Y., Duraisingham, S., Strobel, F. H., Khan, N., Soltow, Q. A., et al. (2013). Predicting network activity from high throughput metabolomics. PLoS Comput Biol 9, e1003123.

Liu, Y., Chen, H., Chen, D., Feng, L., and Zhang, J. (2020). Gut Dysbacteriosis Is Associated with an Imbalanced Cytokines Network in Women with Unexplained Miscarriage. In Review doi:10.21203/rs.3.rs-20415/v1.

Lloyd-Price, J., Mahurkar, A., Rahnavard, G., Crabtree, J., Orvis, J., Hall, A. B., et al. (2017). Strains, functions and dynamics in the expanded Human Microbiome Project. Nature 550, 61–66.

Mardinoglu, A., Shoaie, S., Bergentall, M., Ghaffari, P., Zhang, C., Larsson, E., et al. (2015). The gut microbiota modulates host amino acid and glutathione metabolism in mice. Mol. Syst. Biol. 11, 834. doi:10.15252/msb.20156487.

Nemet, I., Saha, P. P., Gupta, N., Zhu, W., Romano, K. A., Skye, S. M., et al. (2020). A Cardiovascular Disease-Linked Gut Microbial Metabolite Acts via Adrenergic Receptors. Cell 180, 862-877.e22. doi:10.1016/j.cell.2020.02.016.

Norred, W. P., Nishie, K., and Osman, S. F. (1976). Excretion, distribution and metabolic fate of 3H-alpha-chaconine. Res. Commun. Chem. Pathol. Pharmacol. 13, 161–171.

Priyadarshini, M., Thomas, A., Reisetter, A. C., Scholtens, D. M., Wolever, T. M., Josefson, J. L., et al. (2014). Maternal short-chain fatty acids are associated with metabolic parameters in mothers and newborns. Transl. Res. 164, 153–157.

Quinn, R. A., Melnik, A. V., Vrbanac, A., Fu, T., Patras, K. A., Christy, M. P., et al. (2020). Global chemical effects of the microbiome include new bile-acid conjugations. Nature 579, 123–129. doi:10.1038/s41586-020-2047-9.

Raman, S. B., and Rathinasabapathi, B. (2003). β-Alanine N -Methyltransferase of Limonium latifolium. cDNA Cloning and Functional Expression of a Novel N -Methyltransferase Implicated in the Synthesis of the Osmoprotectant β-Alanine Betaine. Plant Physiol. 132, 1642–1651. doi:10.1104/pp.103.020453.

Reigstad, C. S., Salmonson, C. E., Iii, J. F. R., Szurszewski, J. H., Linden, D. R., Sonnenburg, J. L., et al. (2015). Gut microbes promote colonic serotonin production through an effect of shortLJchain fatty acids on enterochromaffin cells. FASEB J. 29, 1395–1403. doi:10.1096/fj.14-259598.

Roediger, W. E. W., and Moore, A. (1981). Effect of short-chain fatty acid on sodium absorption in isolated human colon perfused through the vascular bed. Dig. Dis. Sci. 26, 100–106.

Sender, R., Fuchs, S., and Milo, R. (2016). Are We Really Vastly Outnumbered? Revisiting the Ratio of Bacterial to Host Cells in Humans. Cell 164, 337–340. doi:10.1016/j.cell.2016.01.013.

Servillo, L., D’Onofrio, N., Giovane, A., Casale, R., Cautela, D., Castaldo, D., et al. (2018). Ruminant meat and milk contain δ-valerobetaine, another precursor of trimethylamine N-oxide (TMAO) like γ-butyrobetaine. Food Chem. 260, 193–199. doi:10.1016/j.foodchem.2018.03.114.

Shekhawat, P., Bennett, M. J., Sadovsky, Y., Nelson, D. M., Rakheja, D., and Strauss, A. W. (2003). Human placenta metabolizes fatty acids: implications for fetal fatty acid oxidation disorders and maternal liver diseases. Am. J. Physiol.-Endocrinol. Metab. 284, E1098–E1105.

Smith, P. M., Howitt, M. R., Panikov, N., Michaud, M., Gallini, C. A., Bohlooly-Y, M., et al. (2013). The Microbial Metabolites, Short-Chain Fatty Acids, Regulate Colonic Treg Cell Homeostasis. Science 341, 569–573. doi:10.1126/science.1241165.

Subramanian, A., Tamayo, P., Mootha, V. K., Mukherjee, S., Ebert, B. L., Gillette, M. A., et al. (2005). Gene set enrichment analysis: a knowledge-based approach for interpreting genome-wide expression profiles. Proc. Natl. Acad. Sci. 102, 15545–15550.

Sumner, L. W., Amberg, A., Barrett, D., Beale, M. H., Beger, R., Daykin, C. A., et al. (2007). Proposed minimum reporting standards for chemical analysis: Chemical Analysis Working Group (CAWG) Metabolomics Standards Initiative (MSI). Metabolomics 3, 211–221. doi:10.1007/s11306-007-0082-2.

Swann, J. R., Spitzer, S. O., and Diaz Heijtz, R. (2020). Developmental Signatures of Microbiota-Derived Metabolites in the Mouse Brain. Metabolites 10, 172. doi:10.3390/metabo10050172.

Tang, W. H. W., Wang, Z., Levison, B. S., Koeth, R. A., Britt, E. B., Fu, X., et al. (2013). Intestinal Microbial Metabolism of Phosphatidylcholine and Cardiovascular Risk. http://dx.doi.org/10.1056/NEJMoa1109400. zdoi:10.1056/NEJMoa1109400.

Tsugawa, H., Cajka, T., Kind, T., Ma, Y., Higgins, B., Ikeda, K., et al. (2015). MS-DIAL: data-independent MS/MS deconvolution for comprehensive metabolome analysis. Nat. Methods 12, 523–526. doi:10.1038/nmeth.3393.

van de Pavert, S. A., Ferreira, M., Domingues, R. G., Ribeiro, H., Molenaar, R., Moreira-Santos, L., et al. (2014). Maternal retinoids control type 3 innate lymphoid cells and set the offspring immunity. Nature 508, 123–127. doi:10.1038/nature13158.

Velasquez, M., Ramezani, A., Manal, A., and Raj, D. (2016). Trimethylamine N-Oxide: The Good, the Bad and the Unknown. Toxins 8, 326. doi:10.3390/toxins8110326.

Venkatesh, M., Mukherjee, S., Wang, H., Li, H., Sun, K., Benechet, A. P., et al. (2014). Symbiotic Bacterial Metabolites Regulate Gastrointestinal Barrier Function via the Xenobiotic Sensor PXR and Toll-like Receptor 4. Immunity 41, 296–310. doi:10.1016/j.immuni.2014.06.014.

Vuong, H. E., Pronovost, G. N., Williams, D. W., Coley, E. J. L., Siegler, E. L., Qiu, A., et al. (2020). The maternal microbiome modulates fetal neurodevelopment in mice. Nature. doi:10.1038/s41586-020-2745-3.

Walker, R. W., Clemente, J. C., Peter, I., and Loos, R. J. F. (2017). The prenatal gut microbiome: are we colonized with bacteria in utero? Pediatr. Obes. 12 Suppl 1, 3–17. doi:10.1111/ijpo.12217.

Wang, Z., Klipfell, E., Bennett, B. J., Koeth, R., Levison, B. S., DuGar, B., et al. (2011). Gut flora metabolism of phosphatidylcholine promotes cardiovascular disease. Nature 472, 57–63. doi:10.1038/nature09922.

Wikoff, W. R., Anfora, A. T., Liu, J., Schultz, P. G., Lesley, S. A., Peters, E. C., et al. (2009). Metabolomics analysis reveals large effects of gut microflora on mammalian blood metabolites. Proc. Natl. Acad. Sci. U. S. A. 106, 3698–703. doi:10.1073/pnas.0812874106.

Yamamoto, Y., Nakanishi, Y., Murakami, S., Aw, W., Tsukimi, T., Nozu, R., et al. (2018). A Metabolomic-Based Evaluation of the Role of Commensal Microbiota throughout the Gastrointestinal Tract in Mice. Microorganisms 6, 101. doi:10.3390/microorganisms6040101.

Yano, J. M., Yu, K., Donaldson, G. P., Shastri, G. G., Ann, P., Ma, L., et al. (2015). Indigenous Bacteria from the Gut Microbiota Regulate Host Serotonin Biosynthesis. Cell 161, 264– 276. doi:10.1016/j.cell.2015.02.047.

Yu, Y., Raka, F., and Adeli, K. (2019). The Role of the Gut Microbiota in Lipid and Lipoprotein Metabolism. J. Clin. Med. 8. doi:10.3390/jcm8122227.

Zelante, T., Iannitti, R. G., Cunha, C., De Luca, A., Giovannini, G., Pieraccini, G., et al. (2013). Tryptophan Catabolites from Microbiota Engage Aryl Hydrocarbon Receptor and Balance Mucosal Reactivity via Interleukin-22. Immunity 39, 372–385. doi:10.1016/j.immuni.2013.08.003.

Zhang, L. S., and Davies, S. S. (2016). Microbial metabolism of dietary components to bioactive metabolites: opportunities for new therapeutic interventions. Genome Med. 8, 46. doi:10.1186/s13073-016-0296-x.

Zhu, W., Gregory, J. C., Org, E., Buffa, J. A., Gupta, N., Wang, Z., et al. (2016). Gut Microbial Metabolite TMAO Enhances Platelet Hyperreactivity and Thrombosis Risk. Cell 165, 111–124. doi:10.1016/j.cell.2016.02.011.

